# BMPR2 inhibits activin- and BMP-signaling via wild type ALK2

**DOI:** 10.1101/222406

**Authors:** Oddrun Elise Olsen, Meenu Sankar, Samah Elsaadi, Hanne Hella, Glenn Buene, Sagar Ramesh Darvekar, Kristine Misund, Takenobu Katagiri, Toril Holien

## Abstract

Activin A is a member of the TGF-β superfamily and activates the transcription factors SMAD2/3 through the ALK4 type 1 receptor. Activin A has also been shown to activate SMAD1/5/8 through mutated variants of the type 1 receptor ALK2. Interestingly, we here show that both activin A and activin B could activate SMAD1/5/8 through endogenous wild type ALK2 in multiple myeloma cells. Knockdown of the type 2 receptor BMPR2 strongly potentiated activin A- and activin B-induced SMAD1/5/8 activation and subsequent cell death. Furthermore, activity of BMP6, BMP7 or BMP9, which also signal via ALK2, was potentiated by BMPR2 knockdown. Similar results were seen in HepG2 liver carcinoma cells. We propose that BMPR2 inhibited ALK2-mediated signaling by preventing ALK2 from oligomerizing with the type 2 receptors ACVR2A and ACVR2B, necessary for ALK2 activation by activins and several BMPs in these cells. In conclusion, BMPR2 could be explored as a possible target for therapy in patients with multiple myeloma.

**Summary Statement:** The activation of SMAD1/5/8 via endogenous wild type ALK2 by activin A, activin B, and certain BMPs was enhanced when BMPR2 levels were knocked down.

## Introduction

The transforming growth factor (TGF)-β superfamily of ligands consists of over 30 human members, including bone morphogenetic proteins (BMPs), growth differentiation factors (GDFs), activins and TGF-β isoforms (Wakefield and Hill, 2013). These structurally related ligands are involved in a plethora of processes in the developing and adult organism. The ligands signal by binding to type 1 and type 2 receptors, causing phosphorylation of receptor-activated SMAD transcription factors. In the case of activins, two type 1 receptors are known to relay signals between ligands and SMADs, namely ALK4 (*ACVR1B*) and ALK7 (*ACVR1C*) (ten Dijke et al., 1994; Tsuchida et al., 2004). For BMPs, four different type 1 signaling receptors have been indicated; ALK1 (*ACVRL1*), which primarily is expressed by endothelial cells, and the more ubiquitously expressed ALK2 (*ACVR1*), ALK3 (*BMPR1A*) and ALK6 (*BMPR1B*) (for review on BMP-receptors: (Yadin et al., 2016)). The activin type 1 receptors activate the SMAD2/3-pathway, as TGF-β does through ALK5 (*TGFBR1*), whereas the BMP type 1 receptors activate the SMAD1/5/8-pathway. Two type 2 receptors, ACVR2A and ACVR2B, are shared between activins, BMPs and GDFs, whereas the type 2 receptor BMPR2 only is used by BMPs and GDFs (Yadin et al., 2016).

Activin A has been shown to inhibit growth of normal B cells as well as murine myeloma and plasmacytoma cells, however the mechanism behind was not clear (Brosh et al., 1995; Hashimoto et al., 1998; Nishihara et al., 1993; Zipori and Barda-Saad, 2001). We have earlier shown that activation of SMAD1/5/8 is necessary and sufficient to induce apoptosis in myeloma cells (Holien et al., 2012a). BMP-induced activation of SMAD1/5/8 cause apoptosis in multiple myeloma cells through downregulation of the oncogene c-MYC, which is important for myeloma cell survival (Holien and Sundan, 2014; Holien et al., 2012a; Holien et al., 2012b). Ligands that activate the SMAD2/3-, but not the SMAD1/5/8-pathway, such as TGF-β, do not induce apoptosis in myeloma cells (Baughn et al., 2009; Olsen et al., 2015). We and others have shown that activin A may compete with BMPs for binding to ACVR2A, ACVR2B and ALK2 (Aykul and Martinez-Hackert, 2016; Hatsell et al., 2015; Hino et al., 2015; Olsen et al., 2015; Piek et al., 1999; Seher et al., 2017). Activin A could also induce a weak phosphorylation of SMAD1/5/8 in addition to the canonical activation of SMAD2 (Olsen et al., 2015). This coincided with a modest reduction in myeloma cell viability. Interestingly, it was shown that activin A bind and signal through a mutated variant of ALK2, encoded by *ACVR1* R206H (Hatsell et al., 2015; Hino et al., 2015).

In this study, we wanted to clarify how activins may affect myeloma cell survival. We found that both activin A and activin B signaled through endogenous levels of wild type ALK2 to phosphorylate the BMP-activated SMADs, SMAD1 and SMAD5, and consequently induced myeloma cell death. Surprisingly, depletion of BMPR2 strongly augmented ALK2-, but not ALK4-mediated signaling, induced by either activins or BMPs. In conclusion, we propose that BMPR2 prevents ALK2 from forming active receptor signaling complexes with ACVR2A and ACVR2B, thus inhibiting ALK2-mediated SMAD1/5/8-activation and myeloma cell death.

## Results

### Activin A and activin B inhibited myeloma cells through ALK2 and SMAD1/5/8

Five different myeloma cell lines were treated with increasing doses of activin A or activin B for three days. A dose-dependent decrease in relative viable cell numbers was seen in IH-1 and to a lesser extent in the INA-6 cell line (Fig.1A,B). The other three cell lines were for a large part refractory to activin A- or activin B-induced growth inhibition in the doses used here. Notably, both activins induced canonical activation of SMAD2 in all cell lines, whereas a clear activation of SMAD1/5/8 was also observed in the IH-1, INA-6 and KJON cell lines (Fig. 1C). Contamination by other ligands is a potential issue when using recombinant proteins (Carthy et al., 2016; Olsen et al., 2017). Thus, to confirm that the observed effects were actually caused by activin A and activin B, we performed experiments on SMAD-activity in the presence of neutralizing antibodies (Fig. S1). As shown, antibodies towards activin A and activin B reduced activin A- and activin B-induced SMAD1/5/8 phosphorylation to background levels, indicating that the observed SMAD1/5/8 phosphorylation indeed was due to activin A and activin B and not to contamination factors. We and others have previously shown that activation of SMAD1/5/8, but not SMAD2, induced growth arrest or apoptosis in human myeloma cells (Baughn et al., 2009; Holien et al., 2012a; Olsen et al., 2015). In line with this, we also here show that activin A and activin B induced apoptosis in IH-1 cells as measured by annexin V labeling and cleavage of caspase 3 (Fig. 1D,E). Activin A has previously been shown to signal through ALK4 whereas activin B bind and signal through either ALK4 or ALK7 (Wakefield and Hill, 2013). Neither ALK4 nor ALK7 are thought to activate SMAD1/5/8. Thus, the results above suggest that activin A and activin B also may bind and signal through complexes containing BMP-type 1 receptors.

**Fig. 1.**
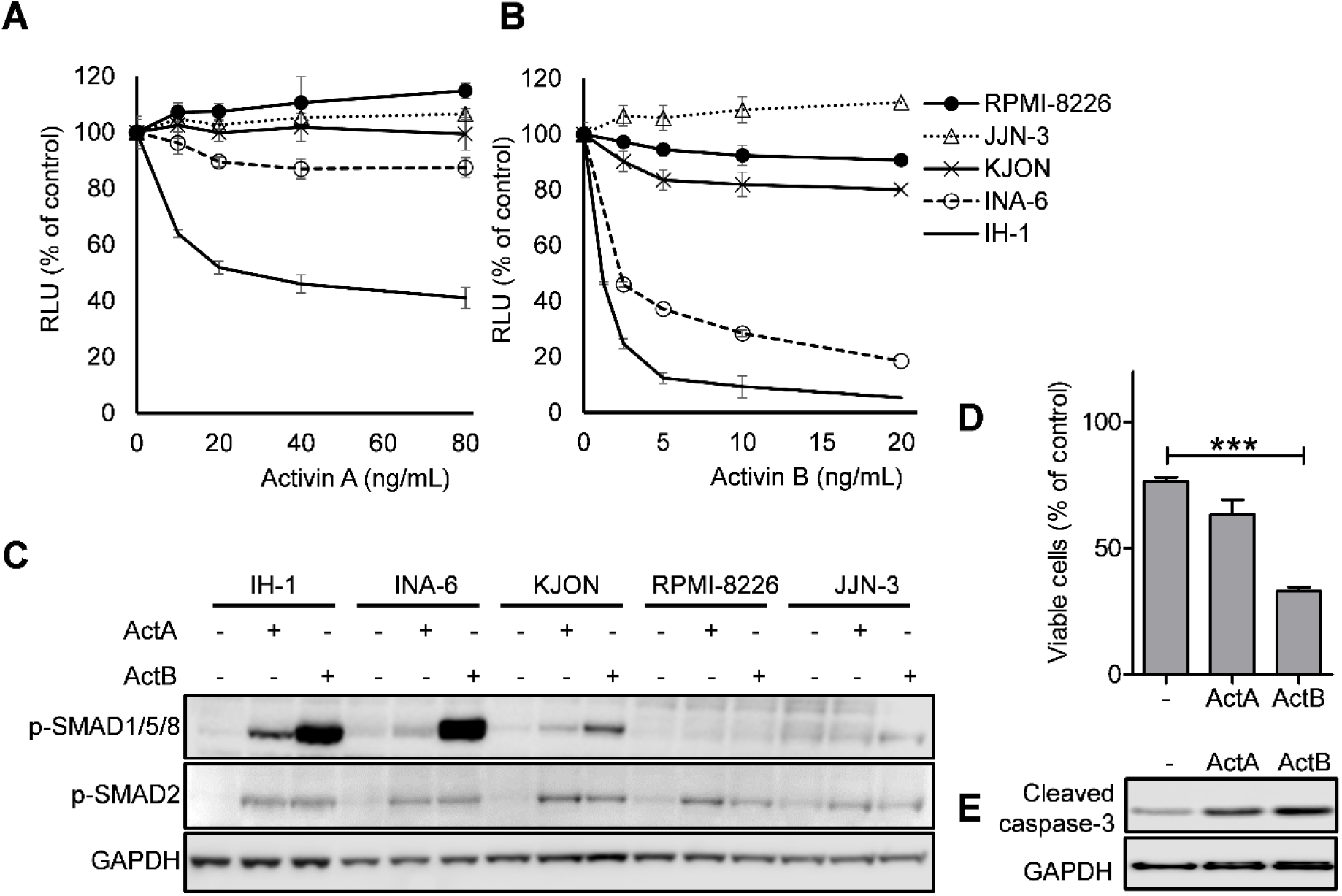
Activin A and activin B inhibited myeloma cells through ALK2 and SMAD1/5/8. The five myeloma cell lines KJON, INA-6, JJN3, RPMI-8226 and IH-1 were treated with increasing doses of activin A (A) or activin B (B) for three days and relative cell numbers were determined using the CellTiter Glo viability assay and expressed as relative luminescence units (RLU). Data are presented as one representative experiment out of three with mean±s.d. of n=3 technical replicates. (C) The cell lines were incubated with activin A or activin B (both 10 ng/mL) for four hours and the cells were subject to immunoblotting with the indicated antibodies. GAPDH was used as loading control. (D) IH-1 cells were treated with activin A (20 ng/mL) or activin B (4 ng/mL) for three days prior to labeling with annexin V-FITC and propidium iodide. The cells that were negative for both were considered viable and plotted in the graph. The graph represents mean±s.e.m. of n=3 independent experiments. A one-way ANOVA, Dunnett’s multiple comparisons test was performed and the number of asterisks displayed over the graph bars reflects on the degree of significance (P ≤ 0.001: ***). (E) IH-1 cells were treated with activin A (20 ng/mL) or activin B (4 ng/mL) overnight and used for immunoblotting with antibodies towards cleaved caspase-3 and GAPDH as a loading control.

### Activins signaled through endogenous wild type ALK2 in myeloma cells

To clarify which BMP-type 1 receptor could be responsible for activin-induced activation of SMAD1/5/8, we first measured the mRNA levels of the potential TGF-β superfamily receptors (Fig. 2A). ALK1/*ACVRL1* and ALK6/*BMPR1B* were also measured, but omitted in the figure since none of the cell lines showed expression of these two receptors. Since the only BMP-type 1 receptor expressed in INA-6 cells was ALK2, whereas IH-1 cells expressed both ALK2 and ALK3, ALK2 was the likely responsible type 1 receptor present in both cell lines. We then employed DMH1 and K02288, inhibitors of SMAD1/5/8-activation, to see their effects on apoptosis induced by activin A and activin B. DMH1 targets BMP-activated ALKs selectively, whereas K02288 has been described as an inhibitor that targets ALK2 more potently than the other BMP-activated ALKs (Hao et al., 2010; Sanvitale et al., 2013). Both inhibitors significantly inhibited the reduction in cell viability caused by activin A and activin B (Fig. 2B,C). We then compared these results with inhibition of ALK4/5/7 by using the SB431542 inhibitor (Inman et al., 2002). This inhibitor showed no significant effect on activin A-induced apoptosis and a modest, but significant effect on activin B-induced apoptosis (Fig. 2D). To target ALK2 even more specifically, we utilized a novel, monoclonal ALK2 neutralizing antibody termed DaVinci (Katagiri et al., 2017). The antibody blunted apoptosis induced by activin A and activin B, as well as by BMP9, which we have previously shown to use ALK2 to activate the SMAD1/5/8-pathway in these cells (Olsen et al., 2014). Taken together, the results support that ALK2 is the BMP-type 1 receptor for activin A and activin B in these cells. Of note, whole exome sequencing of 69 human myeloma cell lines, including INA-6, did not reveal any mutations in ALK2 (data not shown; publicly available data from the Keats lab website; http://www.keatslab.org/data-repository). We also RNA-sequenced our own batch of INA-6 cells. Two single nucleotide variations (SNVs) were detected in the ALK2 gene, *ACVR1*, denoted rs2227861 and rs1146031, none of which cause a change in the amino acid sequence (data not shown). Thus, this we here report for the first time that, not only activin B, but also activin A, can signal through endogenous levels of wild type ALK2.

**Fig. 2.**
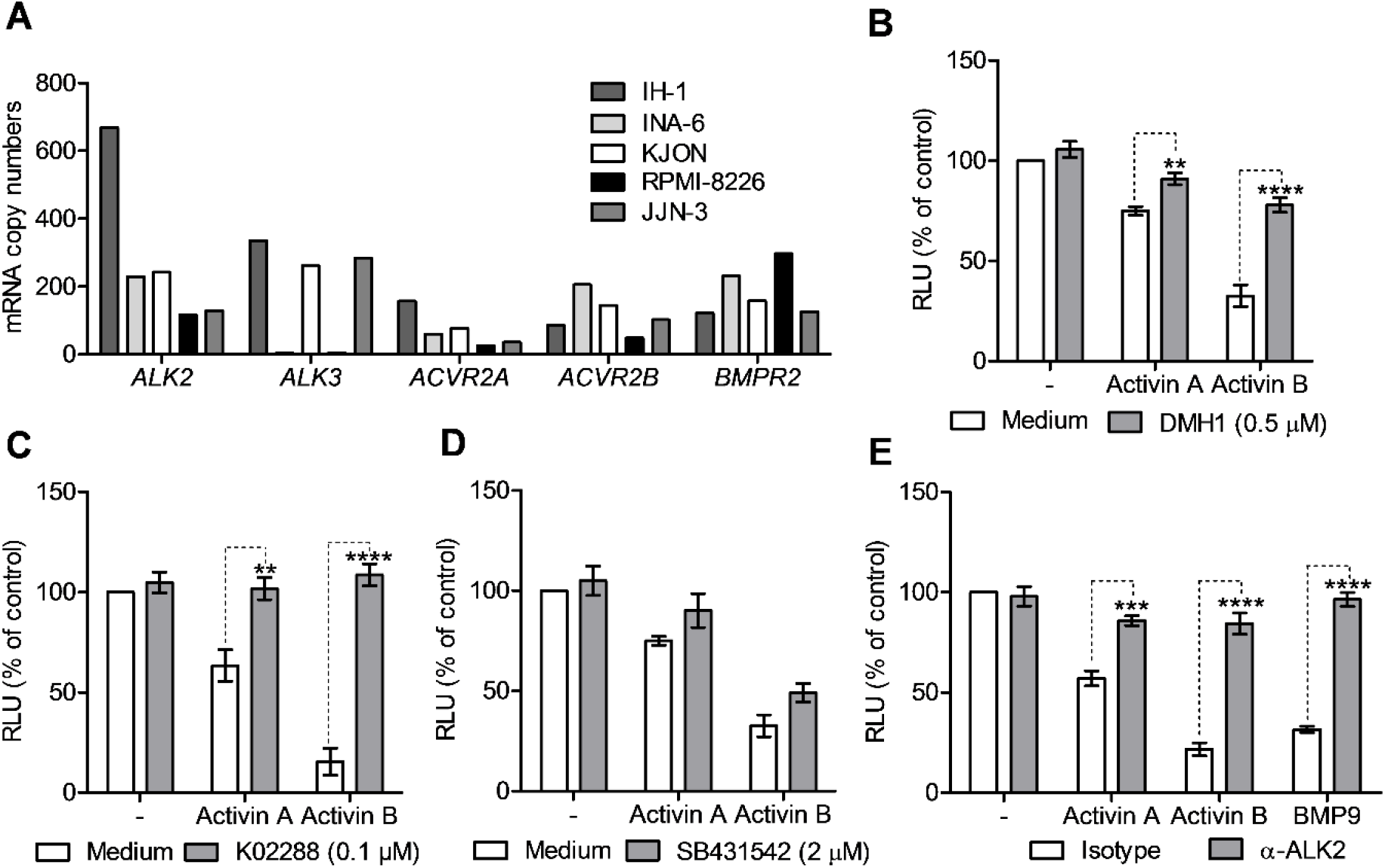
Activins signaled through wild type ALK2 in myeloma cells. (A) The myeloma cell lines used in this study were analyzed by nCounter gene expression analysis for expression of different receptors in the TGF-β superfamily. Shown are values normalized against a set of house-keeping genes and positive controls. ALK1/*ACVRL1* and ALK6/*BMPR1B* were also included in the analysis, but omitted in the plot due to absence of expression in all five cell lines. IH-1 cells were treated with activin A (20 ng/mL) or activin B (4 ng/mL) in the presence of DMH1 (B), K02288 (C), or SB431542 (D), for three days and relative cell numbers were determined using the CellTiter Glo viability assay. (E) IH-1 cells were treated for three days with activin A (40 ng/mL), activin B (1 ng/mL), or BMP9 (0.3 ng/mL) in the presence of an isotype antibody (CTR) or the neutralizing ALK2 antibody, DaVinci (both 10 ng/mL). Relative cell numbers were determined using the CellTiter Glo viability assay. Graphs B-E represent mean±s.e.m. of n=3 independent experiments. Two-way ANOVA, Bonferroni’s multiple comparisons tests were performed and the number of asterisks displayed over the graph bars reflects on the degree of significance (P<0.01: **; P<0.001: ***; P<0.0001: ****). RLU; relative luminescence units.

### Loss of BMPR2 potentiated activin-induced SMAD1/5/8 activity

ACVR2A and ACVR2B are known type 2 receptors for activins and BMPs. Also, BMPR2 is an important BMP type 2 receptor, but the role for BMPR2 in activin-mediated signaling is less clear. We thus generated cells with stable expression of shRNA targeting BMPR2 (shBMPR2), using as controls non-targeting shRNA (shCTR) or shRNA targeting ACVRL1 (shALK1), which is not expressed by myeloma cell lines. Expression of *BMPR2* mRNA was measured in shRNA-expressing cells and indicated a reduction to approximately 10 % and 40 % compared to the levels in control-transduced INA-6 and IH-1 cells, respectively (Fig. 3A,B). BMPR2 protein levels could be detected by immunoblotting in INA-6 cells and there was a clear downregulation of BMPR2 in the shBMPR2-cells compared with control cells (Fig. S2). In IH-1 cells we were not able to detect BMPR2 protein clearly, and thus, we could not determine the degree of downregulation at the protein level. Importantly, activin A activated SMAD1/5/8 more strongly in cells where BMPR2 was downregulated than in control cells, whereas activin A-induced activation of SMAD2 in the same cells was not affected by BMPR2 knockdown (Fig. 3C,D). Also, the mRNA levels of *ID1*, a known SMAD1/5/8 target gene,(Chen et al., 2006) was more potently induced by activin A in the shBMPR2 cells (Fig. 3E,F). Finally, activin A- and activin B-induced growth inhibition was potentiated by knockdown of BMPR2 (Fig. 3G-J). Thus, we here show that activin A- and activin B-induced signaling through ALK2 was potentiated by stable knockdown of BMPR2 both in short- and long-term experiments. The results indicate that presence of BMPR2 inhibits activin-induced ALK2 signaling.

**Fig. 3.**
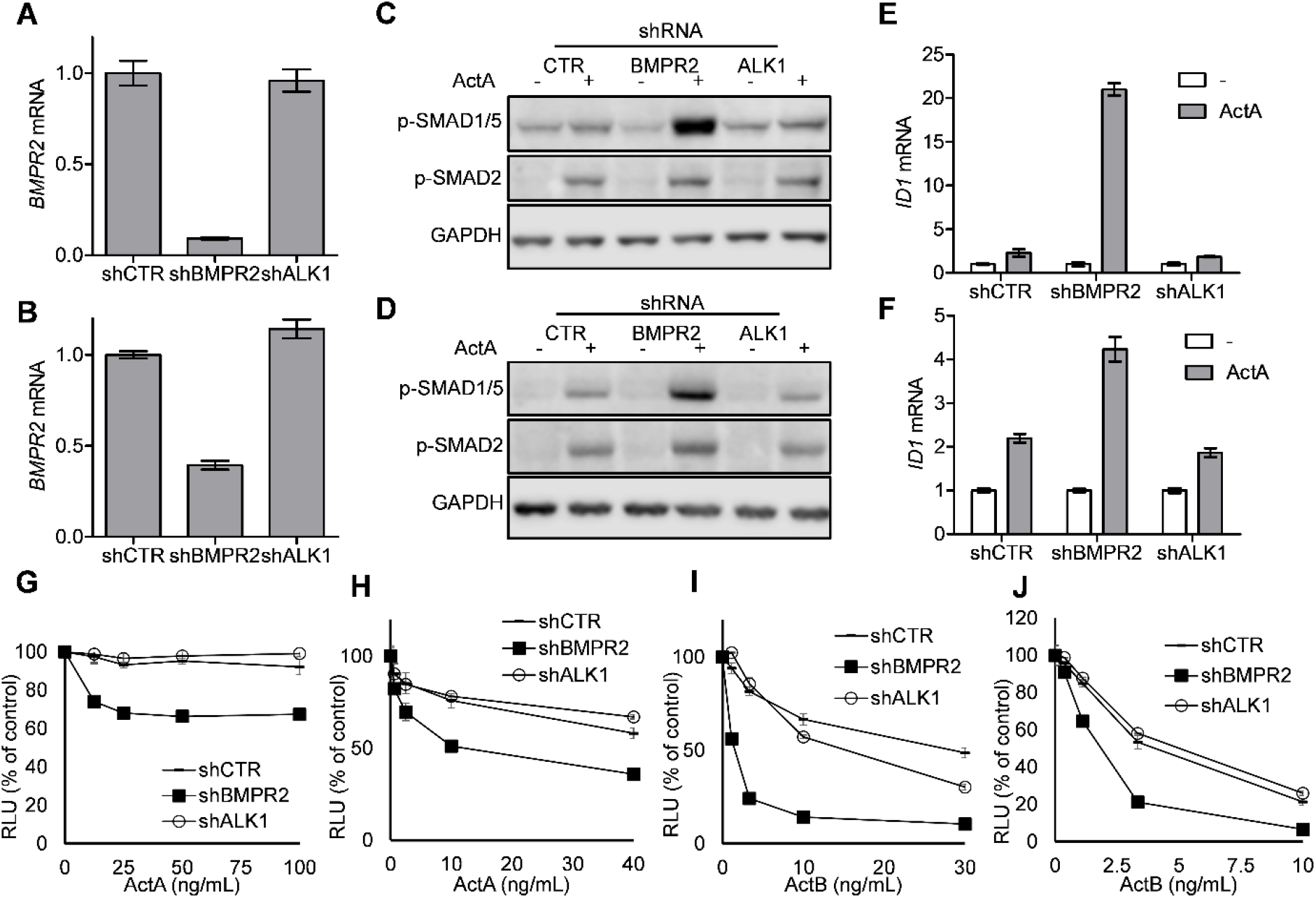
Loss of BMPR2 potentiated activin-induced ALK2 activity. Myeloma cell lines stably expressing shRNAs, shCTR (non-targeting), shBMPR2 or shALK1, were generated. Expression of *BMPR2* mRNA relative to *GAPDH* in the cell lines were determined in INA-6 (A) and IH-1 (B) using the comparative Ct-method. The shRNA-expressing INA-6 cells (C) or IH-1 cells (D) were treated with or without activin A (10 ng/mL) for four hours and used for immunoblotting with the indicated antibodies. INA-6 cells (E) or IH-1 cells (F) were treated with or without activin A (10 ng/mL) for four hours and the expression levels of *ID1* mRNA relative to *GAPDH* were determined by PCR and the comparative Ct-method. INA-6 (G, I) and IH-1 (H, J) cells were treated with increasing doses of activin A or activin B for three days and relative cell numbers were determined using the CellTiter Glo viability assay. Shown are representative experiments of at least three independent experiments with mean±s.d. of n=3 technical replicates. RLU; relative luminescence units.

### Loss of BMPR2 also potentiated the effect of some BMPs

Based on the effects obtained with activins, we wondered if BMP signaling also could be potentiated in the BMPR2 knockdown-cells. The INA-6 cell line expressed the type 2 receptors BMPR2, ACVR2A, and ACVR2B, but only ALK2 as a BMP type 1 receptor (Fig. 2A), and BMP6, BMP7, and BMP9 are the only BMPs currently known to activate SMAD1/5/8 and induce apoptosis in this cell line. IH-1 cells also expressed ALK3 and responds to all BMPs tested by us (BMP2, BMP4, BMP5, BMP6, BMP7, and BMP9) (Hjertner et al., 2001; Olsen et al., 2014; Ro et al., 2004). Both INA-6 (Fig. 4A-C) and IH-1 (Fig. 4D-E) expressing shBMPR2 showed a more pronounced, dose-dependent decrease in viability after treatment with BMP6, BMP7 (only shown for INA-6) or BMP9 than control cells (Fig. 4A-E). In contrast, neither BMP2, BMP4 nor BMP10 treatment was affected by shBMPR2, indicating that these ligands depend on other receptor combinations to signal (Fig. 4F-H).

**Fig. 4.**
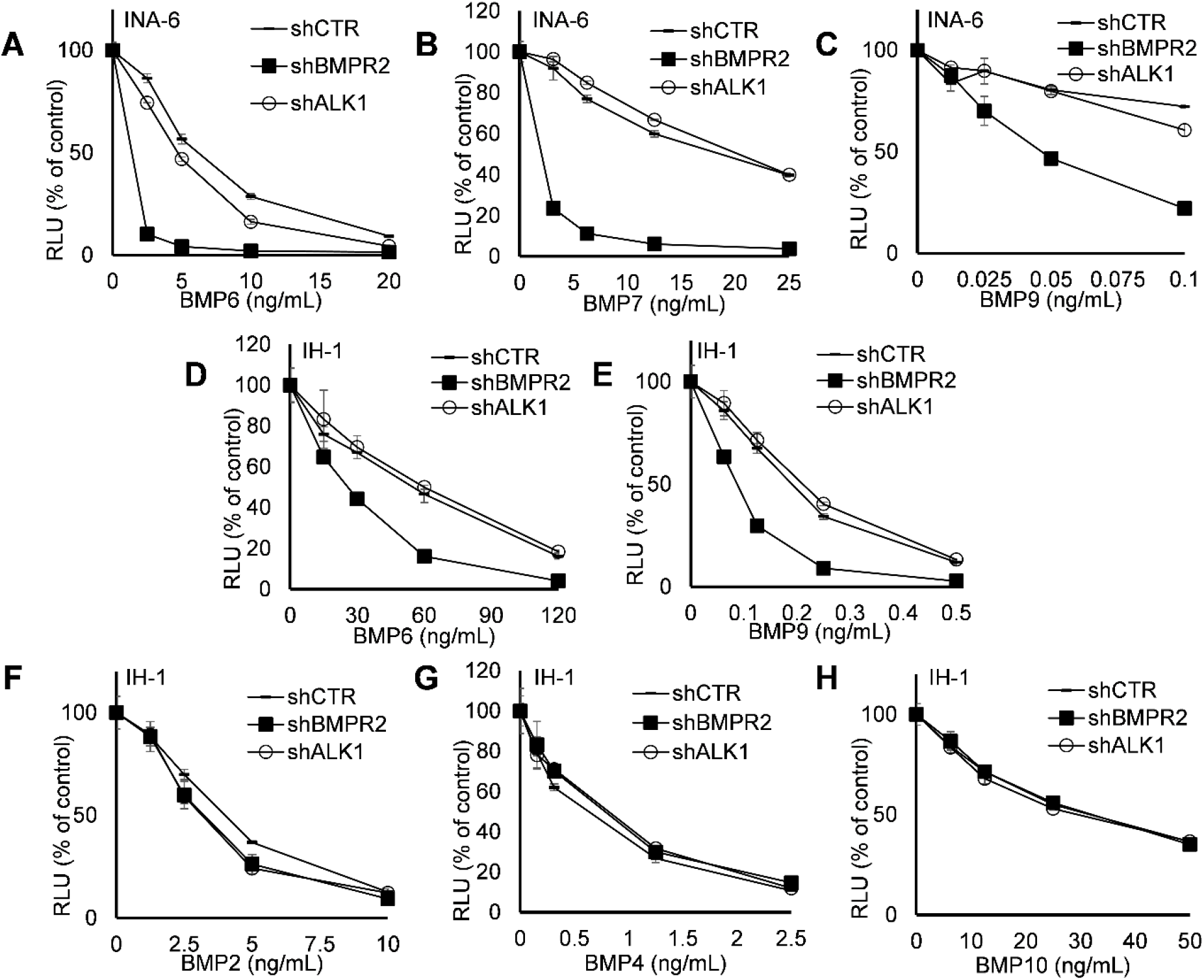
Loss of BMPR2 potentiated the effect of some BMPs. (A-C) Stably expressing INA-6 shCTR, shBMPR2 and shALK1 cells were treated with increasing doses of BMP6, BMP7 or BMP9 for three days and relative cell numbers were determined using the CellTiter Glo assay. (D-H) IH-1 shCTR, shBMPR2 and shALK1 cells were treated with increasing doses of BMP6, BMP9, BMP2, BMP4 or BMP10 for three days and the relative cell numbers were determined by the CellTiter Glo assay. Shown are representative experiments of at least three independent experiments with mean±s.d. of n=3 technical replicates. RLU; relative luminescence units.

### Effect of BMPR2 knockdown in HepG2 liver carcinoma cells

To investigate if the effects of BMPR2 expression levels on BMP- and activin-induced activation of SMAD1/5/8 signaling also could be seen in other cells, we transiently transfected HepG2 liver carcinoma cells with siRNA targeting BMPR2 (siBMPR2) or non-targeting siRNA (siCTR). The cells were treated for four hours with BMP6, activin A or activin B (all at a concentration of 10 ng/mL) and activation of SMADs was determined by immunoblotting. Interestingly, also in HepG2 cells activation of SMAD1/5/8 by these ligands was potentiated by BMPR2 knockdown, whereas activation of SMAD2 remained unaffected (Fig. 5A). The mRNA expression of the SMAD target gene *ID1* in activin A-treated HepG2 cells after BMPR2 knockdown was also augmented (Fig. 5B). The levels of *BMPR2* mRNA in these experiments were reduced to < 20 % of mRNA levels in control cells (Fig. 5C). In summary, also in HepG2 cells there was a potentiation of ligand-induced activation of SMAD1/5/8 by lowering the expression of BMPR2.

**Fig. 5.**
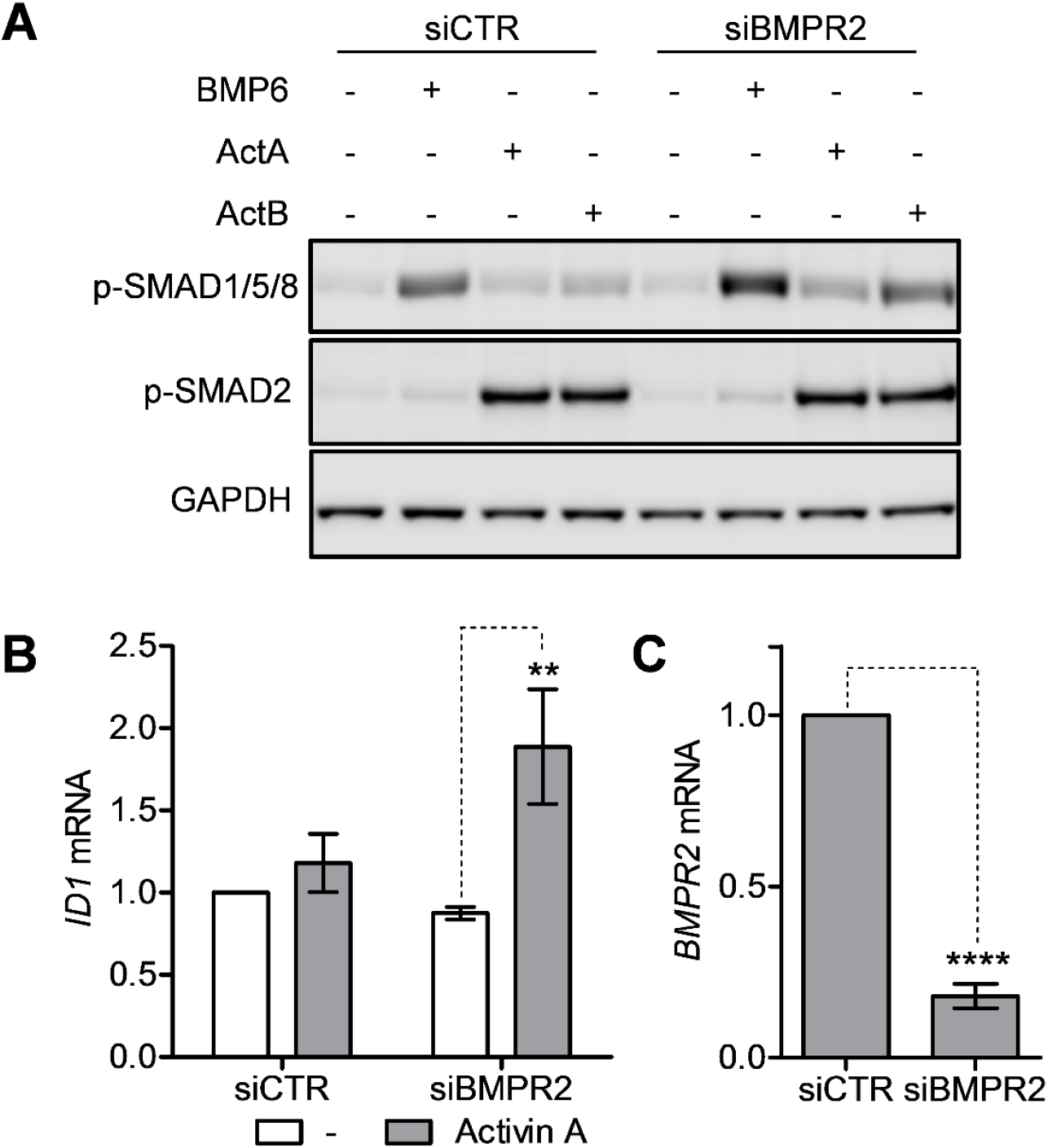
Effect of BMPR2 knockdown in HepG2 liver carcinoma cells. HepG2 liver carcinoma cells were transiently transfected with siRNA targeting BMPR2 (siBMPR2) or non-targeting control (siCTR) and all experiments were performed the day after transfection. (A) Cells were treated with BMP6, activin A or activin B (all at 10 ng/mL) for four hours and subjected to immunoblotting for phospho-SMAD1/5/8, phospho-SMAD2 and GAPDH. Shown is one representative out of three independent experiments. (B) Cells were treated with or without activin A (10 ng/mL) for four hours and expression of *ID1* mRNA relative to *GAPDH* was determined by PCR using the comparative Ct-method. The graph shows mean±s.e.m. of n=4 independent experiments. Two-way ANOVA, Bonferroni’s multiple comparisons test was performed and the number of asterisks displayed over the graph bars reflects on the degree of significance (P<0.01: **). (C) *BMPR2* mRNA levels were measured by PCR using the comparative Ct-method and *GAPDH* as housekeeping gene. The graph shows the mean±s.e.m. of n=4 independent experiments. The number of asterisks displayed over the graph bars reflects on the degree of significance (P<0.0001: ****, one-sided, paired t-test).

## Discussion

The aim of this study was to clarify how activins affected myeloma cell survival. We earlier found that activin A could inhibit BMP6- and BMP9-signaling through ALK2 (Olsen et al., 2015). However, other studies have earlier reported that activin A could induce apoptosis in B cells and murine myeloma and plasmacytoma cells (Brosh et al., 1995; Hashimoto et al., 1998; Nishihara et al., 1993; Zipori and Barda-Saad, 2001). In myeloma cells, activation of the SMAD2 pathway has not been found to induce apoptosis, whereas we earlier found that activation of the SMAD1/5/8-pathway was necessary and sufficient for induction of apoptosis. The most important finding presented here is that activin A and activin B activated the non-canonical SMAD1/5/8 pathway through endogenous wild type ALK2, thus inducing apoptosis in multiple myeloma cells. Moreover, reducing the levels of the type 2 receptor BMPR2 potentiated activation of SMAD1/5/8 through ALK2 by activins as well as by some BMPs that signal through ALK2.

ALK2 was initially described as a type 1 receptor that could bind activin A, but no evidence was found for activin A-induced activation of the SMAD1/5/8 pathway. (Attisano et al., 1993; Ebner et al., 1993; ten Dijke et al., 1994). However, activin A was shown to induce non-canonical activation of SMAD1/5/8 by signaling through a mutated form of ALK2, encoded by *ACVR1* R206H (Hatsell et al., 2015; Hino et al., 2015). Just before submission of this manuscript, there was also a report describing that activin A could induce SMAD1/5/8-activity through overexpressed wild type ALK2, strongly supporting our findings (Haupt et al., 2017). Activin B has been shown to activate SMAD1/5/8 in hepatocytes through ALK2, ACVR2A and ACVR2B (Besson-Fournier et al., 2012; Canali et al., 2016). In line with these results, we here show that both activin A and activin B are able to activate SMAD1/5/8 through the endogenous wild type ALK2 receptor in myeloma cells, leading to cell death.

Activin A-induced SMAD1/5/8-activation and consequently myeloma cell death was strongly augmented when levels of BMPR2 were reduced. Furthermore, the effects of activin B, BMP6, BMP7 and BMP9, which also utilize ALK2 as type 1 receptor, were potentiated in the same way. Our results resemble previously described effects in human mesenchymal stem cells, where BMP2 and BMP4 preferred BMPR2 over ACVR2, whereas BMP6 and BMP7 preferred ACVR2 over BMPR2 for signaling (Lavery et al., 2008). Also, in hepatocytes, knockdown of BMPR2 potentiated BMP6- and activin B-induced SMAD1/5/8 activity (Canali et al., 2016). In pulmonary artery smooth muscle cells (PASMC) loss of BMPR2 augmented signaling by BMP6 and BMP7, whereas BMP2 and BMP4 signaling was diminished (Yu et al., 2005). Contrary to the report on PASMC, we observed no change in the activity of BMP2 or BMP4. We speculate that this may be due to compensatory utilization of the type 2 receptors ACVR2A and ACVR2B by BMP2 and BMP4 in the cell types used here. The same might apply for BMP10-induced growth inhibition, which was also not changed in cells with lower levels of BMPR2. On the other hand, another study found that deleting *Bmpr2* in mouse skeletal progenitor cells selectively impaired activin A-induced SMAD2/3 activation, but had no effect on BMP signaling, causing increased bone formation and bone mass (Lowery et al., 2015). We did not see any significant difference in activin A- or activin B-induced activation of SMAD2 in myeloma cells by lowering BMPR2 levels. The differences between our study and the previously reported results may be due to variations in cellular receptor expression.

Based on our results, one might speculate that increasing BMPR2’s ability to interact with ALK2 could dampen ALK2 signaling. Mutations found in ALK2/ACVR1 in the rare inherited disease FOP increase activin A’s ability to signal through ALK2 (Hatsell et al., 2015; Hino et al., 2015). It has been suggested that the effect of activin A signaling through ACVR1 R206H is cell type specific, i.e. activin A-induced SMAD1/5/8-phosphorylation was induced in human induced pluripotent stem cells (hiPSC), but not in hiPSCs derived endothelial cells (iECs) from FOP patients (Barruet et al., 2016). Interestingly, the expression levels of BMPR2 in hiPSCs were lower than in iECs and in light of our findings, this could possibly explain some of the cell specific observations.

A possible mechanism for the potentiating effects of activin A seen after knocking down BMPR2, could be that activin A bind ALK2-BMPR2 or ALK2-ACVR2 equally well, but induce a weaker signal through ALK2-BMPR2. However, cross-linking experiments with radioactive activin A showed binding of activin A to ALK2 when the receptor was co-expressed with ACVR2A or ACVR2B, but not with BMPR2 (Hino et al., 2015). Another study found that overexpression of Müllerian inhibiting substance receptor II (MISRII) could sequester ALK2. Activin A signaling through ALK4 was thereby potentiated suggesting that presence of ALK2 inhibited activin A signaling by blocking ACVR2 receptor binding to ALK4 (Renlund et al., 2007). The latter study also showed that ALK2 could bind ACVR2A in the absence of ligand. Both studies support our hypothesis that BMPR2 might sequester ALK2 and prevent complex formation between ALK2 and ACVR2A or ACVR2B. A proposed model of our hypothesis is provided (Fig. 6).

**Fig. 6.**
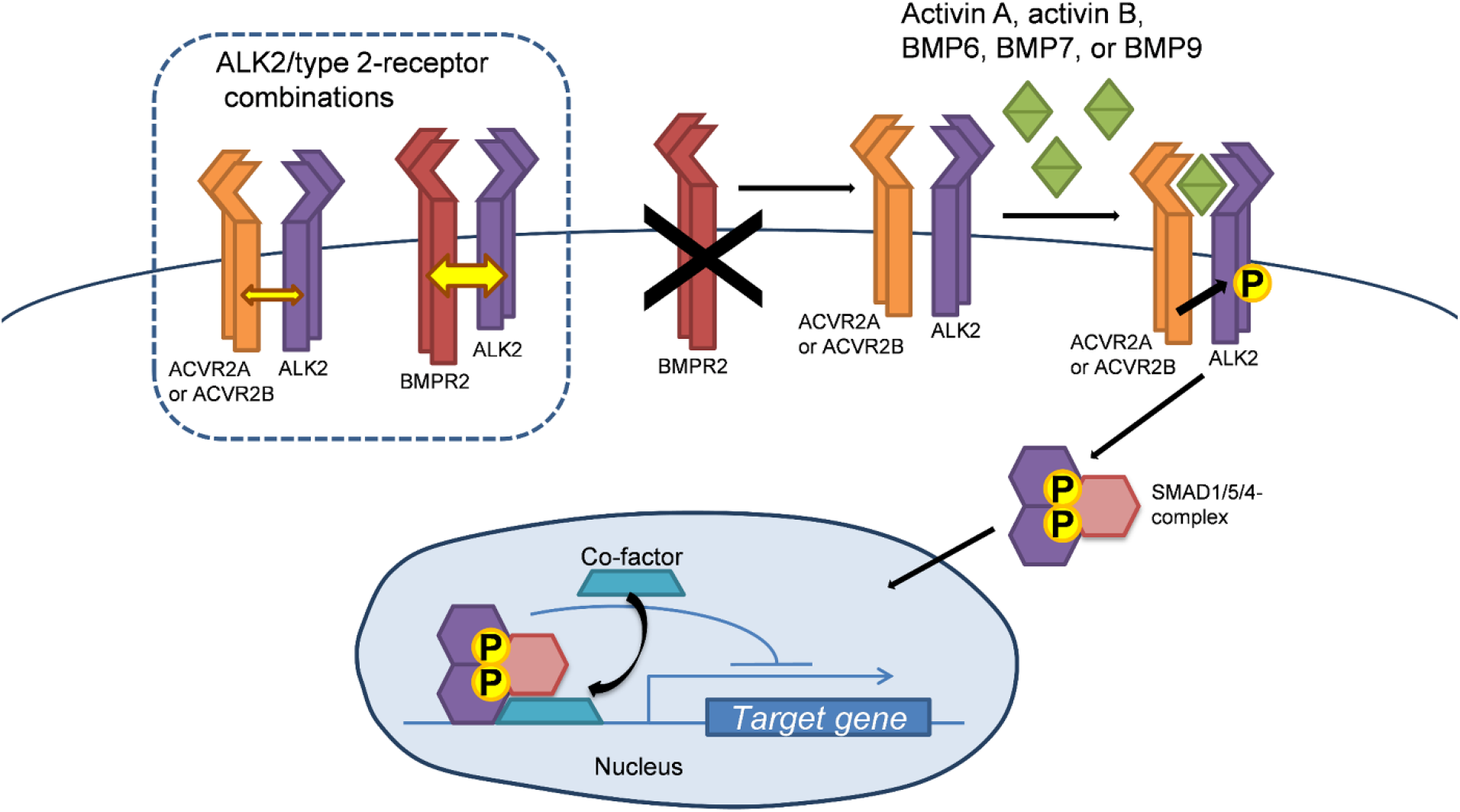
Proposed model of activin- and BMP-signaling through ALK2. In the dotted-line box to the left the possible combinations of ALK2 with a type 2 receptor are shown. Yellow arrows suggest receptor-complex preference. If ALK2 is a limiting factor, the preference for complex formation with a given type 2 receptor will determine signaling outcome. Thus, when levels of BMPR2 are lowered, more ALK2 may be available for complex formation with ACVR2A or ACVR2B, necessary for proper ALK2-signaling by the ligands activin A, activin B, BMP6, BMP7, or BMP9.

This is the first report to show that the TGF-β family ligands activin B and BMP10 can induce apoptosis in myeloma cells. Activin B has been shown to bind ALK2 and activate SMAD1/5/8 in hepatocytes (Canali et al., 2016). Both activin B and BMP10 have been shown to bind with strong affinity to the type 2 receptors ACVR2A and ACVR2B (Aykul and Martinez-Hackert, 2016; Koncarevic et al., 2012). We found that also in myeloma cells, activin B could signal via ALK2 to activate SMAD1/5/8, thus inducing apoptosis. BMP10 is grouped together with BMP9 based on amino acid sequence similarities (Mazerbourg et al., 2005). Like BMP9, BMP10 binds with high affinity to the type 1 receptor ALK1 (David et al., 2007). However, in multiple myeloma cells BMP9 has been shown to use ALK2 (Olsen et al., 2014). Here, BMP10 induced apoptosis in IH-1 cells expressing ALK2 and ALK3. However, INA-6 cells, that only express the BMP type 1 receptor ALK2, were not affected by BMP10 treatment (data not shown). Further studies are needed to determine the exact involvement of type 1 receptors for BMP10 in myeloma cells.

BMPs and activins can be targeted clinically by use of soluble decoy receptors (for a recent review, see (Lowery et al., 2016)). This approach is based on the ligands’ affinities to receptor and may not be a specific way of targeting a particular signaling pathway. In attempts to treat multiple myeloma, it may be better to increase the BMP activity specifically in the cancerous cells. Our results indicate that one way of doing this could be by reducing the levels of BMPR2. To conclude, we hypothesize that reducing the levels of BMPR2 enables increased formation of endogenous receptor complexes of ALK2 with ACVR2A or ACVR2B. The BMPR2-levels could thus dictate the degree of SMAD1/5/8-activation by activin A, activin B and several BMPs. Increased BMPR2 expression in myeloma cells could therefore be one way of escaping the tumor suppressive effects of activins and BMPs. Thus, lowering BMPR2 levels in myeloma cells may be beneficial; but this issue needs to be investigated further.

## Materials and methods

### Cells and reagents

The human multiple myeloma cell lines INA-6 and JJN-3 were kind gifts from Dr. M. Gramatzki (University of Erlangen-Nurnberg, Erlangen, Germany) and Dr. J. Ball (University of Birmingham, UK), respectively. RPMI-8226 cells were from American Type Culture Collection (ATCC; Rockville, MD, USA). IH-1 and KJON were established in our laboratory (Hjertner et al., 2001; Vatsveen et al., 2016). The myeloma cell lines were grown as described previously (Holien et al., 2015). The hepatocyte carcinoma cell line HepG2 was from European Collection of Authenticated Cell Cultures (ECACC; Salisbury, UK). HepG2 cells were grown in 10 % FCS in Eagle’s minimum essential medium supplemented with 2 mM glutamine and non-essential amino acids (Sigma-Aldrich Norway, Oslo, Norway). All cell lines were cultured at 37 ^o^C in a humidified atmosphere containing 5 % CO_2_ and tested regularly for mycoplasma. Fingerprinting was used to confirm the authenticity of the myeloma cell lines. For experiments 2 % human serum (HS) (Department of Immunology and Transfusion Medicine, St. Olav’s University Hospital, Trondheim, Norway) in RPMI was used as medium, with IL-6 (1 ng/mL) mL (Gibco, Thermo Fisher Scientific, Waltham, MA, USA) added for INA-6 and KJON. Apart from IL-6, all recombinant human proteins were from R&D Systems (Bio-Techne, Abingdon, UK). The neutralizing rat monoclonal ALK2 antibody, DaVinci, was a kind gift from Saitama Medical University (Saitama, Japan).

### Immunoblotting

Cells treated as indicated were used for immunoblotting as previously described (Olsen et al., 2015). Primary antibodies used were: phospho-SMAD1/5 (RRID: AB_491015, Cat# 9516), phospho-SMAD1/5/9 (RRID: AB_2493181, Cat# 13820), phospho-SMAD2 (RRID: AB_2631089, Cat# 8828, and cleaved caspase-3 (RRID: AB_2070042, Cat# 9664), all from Cell Signaling Technology, Medprobe, Oslo, Norway, and GAPDH (RRID: AB_2107448, Cat# Ab8245, Abcam, Cambridge, UK). Antibodies were diluted 1:1000, except for GAPDH which was diluted 1:30000.

### Cell viability

Relative viable cell numbers were determined by the CellTiter-Glo assay (Promega, Madison, WI, USA), which measures ATP levels, as described (Olsen et al., 2015).

### Apoptosis

To measure changes in cell viability, cells were stained using Apotest FITC kit (Nexins Research, Kattendijke, The Netherlands). In brief, cells were incubated with annexin V FITC (0.2 μg/ mL in 1X binding buffer) for one hour on ice. Propidium iodide (PI) (1.4 μg/mL) was added five minutes prior to data acquisition using an LSRII flow cytometer (BD Biosciences, San Jose, CA). Cells negative for both annexin V and PI staining were considered viable.

### Generation of shRNA-expressing cell lines

ShRNA constructs were purchased as ready-made lentiviral particles and consisted of pools of three different sequences targeting *BMPR2* (sc-40220-V) or *ALK1* (sc-40212-V), and non-targeting control shRNA (sc-108080) (Santa Cruz Biotechnology, Inc., Heidelberg, Germany). Briefly, cells were treated with lentiviral particles at a multiplicity of infection (MOI) of 15 in the presence of polybrene (8 μg/mL). Fresh medium was added after 24 hours. Puromycin was added after 48 hours to select for cells expressing the shRNA and the recovered viable cells were used for further experiments.

### Comparative RT-PCR

RNA-isolation, complementary DNA (cDNA)-synthesis and PCR was performed using StepOne Real-Time PCR System and Taqman Gene Expression Assays (Applied Biosystems, Thermo Fisher Scientific) as described previously (Olsen et al., 2015). The Taqman assays used were: *ACVR1* (Hs00153836_m1), *ACVR2A* (Hs00155658_m1), *ACVR2B* (Hs00609603_m1), *BMPR2* (Hs00176148_m1 and Hs01574531_m1), *GAPDH* (Hs99999905_m1) and *ID1* (Hs00357821_g1). The comparative Ct method was used to calculate relative changes in gene expression with *GAPDH* as housekeeping gene.

### Nanostring Gene Expression Analysis

For mRNA transcript counting we used nCounter Technology (Nanostring Technologies, Seattle, WA, USA) and a custom-made TGF-β nCounter Kit. The standard mRNA gene expression experiment protocol provided by Nanostring was used. Briefly, 100 ng total RNA from myeloma cell lines was hybridized with reporter probes overnight at 65 °C. The nSolver Analysis Software (Nanostring) was used for calculations of transcript numbers. Sample data was normalized against internal kit positive controls and the following housekeeping genes: *ABCF1, EEF1G, GAPDH, OAZ1*, *POLR2A, RPL19*, and *TUBB*.

### Transient knockdown in HepG2

HepG2 cells were seeded in 6-well plates and left over night to adhere. Cells at a confluence of about 50 % were transfected with siRNA using Lipofectamine RNAiMAX (Invitrogen) according to the protocol. For *BMPR2* knockdown, a mix of two different Silencer Select siRNAs was used (siRNA IDs s2045 and s2046) and compared with a mix of Silencer Select Negative Control No. 1 and No. 2 (Ambion, Thermo Fisher Scientific). The day after transfection, the cells were treated with the indicated ligands for four hours and harvested for immunoblotting or PCR.

### Statistical analysis

GraphPad Prism 7 (Graphpad Software, Inc., San Diego, LA) was used to analyze statistical significance. The tests used for each particular experiment is described in the figure legends.

## Acknowledgments

The authors are grateful for skilled technical help from Berit Størdal.

## References

Attisano, L., Carcamo, J., Ventura, F., Weis, F. M., Massague, J. and Wrana, J. L. (1993). Identification of human activin and TGF beta type I receptors that form heteromeric kinase complexes with type II receptors. Cell 75, 671–680.

Aykul, S. and Martinez-Hackert, E. (2016). Transforming Growth Factor-beta Family Ligands Can Function as Antagonists by Competing for Type II Receptor Binding. J Biol Chem 291, 10792–10804.

Barruet, E., Morales, B. M., Lwin, W., White, M. P., Theodoris, C. V., Kim, H., Urrutia, A., Wong, S. A., Srivastava, D. and Hsiao, E. C. (2016). The ACVR1 R206H mutation found in fibrodysplasia ossificans progressiva increases human induced pluripotent stem cell-derived endothelial cell formation and collagen production through BMP-mediated SMAD1/5/8 signaling. Stem Cell Res Ther 7, 115.

Baughn, L. B., Di Liberto, M., Niesvizky, R., Cho, H. J., Jayabalan, D., Lane, J., Liu, F. and Chen-Kiang, S. (2009). CDK2 phosphorylation of Smad2 disrupts TGF-beta transcriptional regulation in resistant primary bone marrow myeloma cells. J Immunol 182, 1810–1817.

Besson-Fournier, C., Latour, C., Kautz, L., Bertrand, J., Ganz, T., Roth, M. P. and Coppin, H. (2012). Induction of activin B by inflammatory stimuli up-regulates expression of the iron-regulatory peptide hepcidin through Smad1/5/8 signaling. Blood 120, 431–439.

Brosh, N., Sternberg, D., Honigwachs-Sha’anani, J., Lee, B. C., Shav-Tal, Y., Tzehoval, E., Shulman, L. M., Toledo, J., Hacham, Y., Carmi, P., et al. (1995). The plasmacytoma growth inhibitor restrictin-P is an antagonist of interleukin 6 and interleukin 11. Identification as a stroma-derived activin A. J Biol Chem 270, 29594–29600.

Canali, S., Core, A. B., Zumbrennen-Bullough, K. B., Merkulova, M., Wang, C. Y., Schneyer, A. L., Pietrangelo, A. and Babitt, J. L. (2016). Activin B Induces Noncanonical SMAD1/5/8 Signaling via BMP Type I Receptors in Hepatocytes: Evidence for a Role in Hepcidin Induction by Inflammation in Male Mice. Endocrinology 157, 1146–1162.

Carthy, J. M., Engstrom, U., Heldin, C. H. and Moustakas, A. (2016). Commercially Available Preparations of Recombinant Wnt3a Contain Non-Wnt Related Activities Which May Activate TGF-beta Signaling. J Cell Biochem 117, 938–945.

Chen, X., Zankl, A., Niroomand, F., Liu, Z., Katus, H. A., Jahn, L. and Tiefenbacher, C. (2006). Upregulation of ID protein by growth and differentiation factor 5 (GDF5) through a smad-dependent and MAPK-independent pathway in HUVSMC. J Mol Cell Cardiol 41, 26–33.

David, L., Mallet, C., Mazerbourg, S., Feige, J. J. and Bailly, S. (2007). Identification of BMP9 and BMP10 as functional activators of the orphan activin receptor-like kinase 1 (ALK1) in endothelial cells. Blood 109, 1953–1961.

Ebner, R., Chen, R. H., Lawler, S., Zioncheck, T. and Derynck, R. (1993). Determination of type I receptor specificity by the type II receptors for TGF-beta or activin. Science 262, 900–902.

Hao, J., Ho, J. N., Lewis, J. A., Karim, K. A., Daniels, R. N., Gentry, P. R., Hopkins, C. R., Lindsley, C. W. and Hong, C. C. (2010). In vivo structure-activity relationship study of dorsomorphin analogues identifies selective VEGF and BMP inhibitors. ACS Chem Biol 5, 245–253.

Hashimoto, O., Yamato, K., Koseki, T., Ohguchi, M., Ishisaki, A., Shoji, H., Nakamura, T., Hayashi, Y., Sugino, H. and Nishihara, T. (1998). The role of activin type I receptors in activin A-induced growth arrest and apoptosis in mouse B-cell hybridoma cells. Cell Signal 10, 743–749.

Hatsell, S. J., Idone, V., Wolken, D. M., Huang, L., Kim, H. J., Wang, L., Wen, X., Nannuru, K. C., Jimenez, J., Xie, L., et al. (2015). ACVR1R206H receptor mutation causes fibrodysplasia ossificans progressiva by imparting responsiveness to activin A. Sci Transl Med 7, 303ra137.

Haupt, J., Xu, M. and Shore, E. M. (2017). Variable signaling activity by FOP ACVR1 mutations. Bone.

Hino, K., Ikeya, M., Horigome, K., Matsumoto, Y., Ebise, H., Nishio, M., Sekiguchi, K., Shibata, M., Nagata, S., Matsuda, S., et al. (2015). Neofunction of ACVR1 in fibrodysplasia ossificans progressiva. Proc Natl Acad Sci U S A 112, 15438–15443.

Hjertner, O., Hjorth-Hansen, H., Borset, M., Seidel, C., Waage, A. and Sundan, A. (2001). Bone morphogenetic protein-4 inhibits proliferation and induces apoptosis of multiple myeloma cells. Blood 97, 516–522.

Holien, T., Misund, K., Olsen, O. E., Baranowska, K. A., Buene, G., Borset, M., Waage, A. and Sundan, A. (2015). MYC amplifications in myeloma cell lines: correlation with MYC-inhibitor efficacy. Oncotarget 6, 22698–22705.

Holien, T. and Sundan, A. (2014). The role of bone morphogenetic proteins in myeloma cell survival. Cytokine Growth Factor Rev 25, 343–350.

Holien, T., Vatsveen, T. K., Hella, H., Rampa, C., Brede, G., Groseth, L. A., Rekvig, M., Borset, M., Standal, T., Waage, A., et al. (2012a). Bone morphogenetic proteins induce apoptosis in multiple myeloma cells by Smad-dependent repression of MYC. Leukemia 26, 1073–1080.

Holien, T., Vatsveen, T. K., Hella, H., Waage, A. and Sundan, A. (2012b). Addiction to c-MYC in multiple myeloma. Blood 120, 2450–2453.

Inman, G. J., Nicolas, F. J., Callahan, J. F., Harling, J. D., Gaster, L. M., Reith, A. D., Laping, N. J. and Hill, C. S. (2002). SB-431542 is a potent and specific inhibitor of transforming growth factor-beta superfamily type I activin receptor-like kinase (ALK) receptors ALK4, ALK5, and ALK7. Mol Pharmacol 62, 65–74.

Katagiri, T., Tsuji, S., Tsukamoto, S., Ohte, S., Kumagai, K., Osawa, K., Takaishi, K., Nakamura, K., Kawaguchi, Y. and Hasegawa, J. (2017). Development of blocking monoclonal antibodies against ALK2, which is a type I receptor for BMPs. J Bone Miner Res 32, Suppl 1, Available at http://www.asbmr.org/education/AbstractDetail?aid=ac10e419-e975-46a4-b6db-d192adf37806. Accessed November 1., 2017.

Koncarevic, A., Kajimura, S., Cornwall-Brady, M., Andreucci, A., Pullen, A., Sako, D., Kumar, R., Grinberg, A. V., Liharska, K., Ucran, J. A., et al. (2012). A novel therapeutic approach to treating obesity through modulation of TGFbeta signaling. Endocrinology 153, 3133–3146.

Lavery, K., Swain, P., Falb, D. and Alaoui-Ismaili, M. H. (2008). BMP-2/4 and BMP-6/7 differentially utilize cell surface receptors to induce osteoblastic differentiation of human bone marrow-derived mesenchymal stem cells. J Biol Chem 283, 20948–20958.

Lowery, J. W., Brookshire, B. and Rosen, V. (2016). A Survey of Strategies to Modulate the Bone Morphogenetic Protein Signaling Pathway: Current and Future Perspectives. Stem Cells Int 2016, 7290686.

Lowery, J. W., Intini, G., Gamer, L., Lotinun, S., Salazar, V. S., Ote, S., Cox, K., Baron, R. and Rosen, V. (2015). Loss of BMPR2 leads to high bone mass due to increased osteoblast activity. J Cell Sci 128, 1308–1315.

Mazerbourg, S., Sangkuhl, K., Luo, C. W., Sudo, S., Klein, C. and Hsueh, A. J. (2005). Identification of receptors and signaling pathways for orphan bone morphogenetic protein/growth differentiation factor ligands based on genomic analyses. J Biol Chem 280, 32122–32132.

Nishihara, T., Okahashi, N. and Ueda, N. (1993). Activin A induces apoptotic cell death. Biochem Biophys Res Commun 197, 985–991.

Olsen, O. E., Skjaervik, A., Stordal, B. F., Sundan, A. and Holien, T. (2017). TGF-β contamination of purified recombinant GDF15. PLoS One 12, e0187349.

Olsen, O. E., Wader, K. F., Hella, H., Mylin, A. K., Turesson, I., Nesthus, I., Waage, A., Sundan, A. and Holien, T. (2015). Activin A inhibits BMP-signaling by binding ACVR2A and ACVR2B. Cell Commun Signal 13, 27.

Olsen, O. E., Wader, K. F., Misund, K., Vatsveen, T. K., Ro, T. B., Mylin, A. K., Turesson, I., Stordal, B. F., Moen, S. H., Standal, T., et al. (2014). Bone morphogenetic protein-9 suppresses growth of myeloma cells by signaling through ALK2 but is inhibited by endoglin. Blood Cancer J 4, e196.

Piek, E., Afrakhte, M., Sampath, K., van Zoelen, E. J., Heldin, C. H. and ten Dijke, P. (1999). Functional antagonism between activin and osteogenic protein-1 in human embryonal carcinoma cells. Journal of cellular physiology 180, 141–149.

Renlund, N., O’Neill, F. H., Zhang, L., Sidis, Y. and Teixeira, J. (2007). Activin receptor-like kinase-2 inhibits activin signaling by blocking the binding of activin to its type II receptor. J Endocrinol 195, 95103.

Ro, T. B., Holt, R. U., Brenne, A. T., Hjorth-Hansen, H., Waage, A., Hjertner, O., Sundan, A. and Borset, M. (2004). Bone morphogenetic protein-5, −6 and −7 inhibit growth and induce apoptosis in human myeloma cells. Oncogene 23, 3024–3032.

Sanvitale, C. E., Kerr, G., Chaikuad, A., Ramel, M. C., Mohedas, A. H., Reichert, S., Wang, Y., Triffitt, J. T., Cuny, G. D., Yu, P. B., et al. (2013). A new class of small molecule inhibitor of BMP signaling. PLoS One 8, e62721.

Seher, A., Lagler, C., Stuhmer, T., Muller-Richter, U. D. A., Kubler, A. C., Sebald, W., Muller, T. D. and Nickel, J. (2017). Utilizing BMP-2 muteins for treatment of multiple myeloma. PLoS One 12, e0174884.

ten Dijke, P., Yamashita, H., Ichijo, H., Franzen, P., Laiho, M., Miyazono, K. and Heldin, C. H. (1994). Characterization of type I receptors for transforming growth factor-beta and activin. Science 264, 101–104.

Tsuchida, K., Nakatani, M., Yamakawa, N., Hashimoto, O., Hasegawa, Y. and Sugino, H. (2004). Activin isoforms signal through type I receptor serine/threonine kinase ALK7. Mol Cell Endocrinol 220, 59–65.

Vatsveen, T. K., Borset, M., Dikic, A., Tian, E., Micci, F., Lid, A. H., Meza-Zepeda, L. A., Coward, E., Waage, A., Sundan, A., et al. (2016). VOLIN and KJON-Two novel hyperdiploid myeloma cell lines. Genes Chromosomes Cancer 55, 890–901.

Wakefield, L. M. and Hill, C. S. (2013). Beyond TGFbeta: roles of other TGFbeta superfamily members in cancer. Nat Rev Cancer 13, 328–341.

Yadin, D., Knaus, P. and Mueller, T. D. (2016). Structural insights into BMP receptors: Specificity, activation and inhibition. Cytokine Growth Factor Rev 27, 13–34.

Yu, P. B., Beppu, H., Kawai, N., Li, E. and Bloch, K. D. (2005). Bone morphogenetic protein (BMP) type II receptor deletion reveals BMP ligand-specific gain of signaling in pulmonary artery smooth muscle cells. J Biol Chem 280, 24443–24450.

Zipori, D. and Barda-Saad, M. (2001). Role of activin A in negative regulation of normal and tumor B lymphocytes. J Leukoc Biol 69, 867–873.

